# Engineered BCL6 BTB Domain of the Bcl-2 Protein Family shows Dynamic Structural Behavior: Insights from Molecular Dynamics Simulations

**DOI:** 10.1101/2023.08.12.553103

**Authors:** Md. Saddam, Mohammad Ahsan Habib, Md. Abrar Fahim, Afsana Mimi, Saiful Islam, Md Mostofa Uddin Helal

## Abstract

Apoptosis is crucially regulated by the Bcl-6 protein, and mutations in this protein can have a significant impact on many malignancies. In this study, we used molecular dynamics simulations to examine the effects of specific mutations (Q8C, R67C, and N84C) in the crystal structure of the BCL6 BTB domain in a compound with pyrazole-pyrimidine ligand. We concentrated on comprehending the dynamics of these alterations and their possible effects on the emergence of cancer. To explore the structural and dynamic changes induced by these mutations, we performed in silico simulations using the GROMACS software suite (version 5.2, 2020.1) on Google Colab’s Tesla T4 GPU. The crystal structure of the BCL6 BTB domain in complex with the pyrazole-pyrimidine ligand (PDB ID: 5N20) served as the wild-type reference structure. Mutations were imposed using the Rotamer functions of Chimera. The simulations were carried out for a total duration of 20 ns using a time step of 2 femtoseconds (0.002 ps). The Trajectory profiles of the BCL6 BTB domain protein and its three mutations, Q8C, R67C and N84C, were shown to differ from each other. Based on the analysis of RMSD, RMSF, and Rg, it was determined that the mutant 2 (R67C) protein exhibited increased instability and greater flexibility. In contrast, mutant 3 (N84C) demonstrates a heightened level of compactness and greater stability compared to the remaining protein mutant. PCA also provides information regarding the structural dynamics of these mutants. In addition, the SASA and SASA autocorrelation provides a distinct view of the solvent accessibility of these proteins.

## 1. Introduction

The nuclear transcriptional repressor that the Bcl-6 proto-oncogene encodes serves a crucial part in the development of germinal centres (GCs) and the control of lymphocyte function, differentiation, and survival[1] that is essential for the start and upkeep of the GC reaction and is a member of the Broad-Complex, Tramtrack, Bric-a-brac/Pox virus and Zinc Finger (BTB/POZ) family[2]. BCL-6 assisted in creating a chromatin accessibility landscape that was favorable for the expression of genes involved in development and hindering for the expression of genes linked to naive T cell programming[3] and protects GC B cells from premature activation and differentiation and creates an environment that is tolerant of the DNA breaks brought on by the remodeling processes of the immunoglobulin gene that result in the creation of high-affinity antibodies of various isotypes[4]. That is commonly translocated in lymphomas, controls inflammation and germinal centre B cell development[5]. Angioimmunoblastic T-cell lymphomas (AITL) and peripheral T-cell lymphomas (PTCL) with TFH-like tumour cells show high levels of BCL6, a master transcription factor for follicular helper T (TFH) cell development[6]. Bcl-6 is required for the growth of several immune system cell types, including germinal center B cells and CD4+ follicular helper T cells[7] the role of these related epigenetic mechanisms in the growth of B-cell lymphoma[8]. TH1, TH2, and TH17 effector cell lineages are not involved in how mature follicular cells support B cells; rather, interleukin-6 and interleukin-21 act as mediators[9].The most often involved oncogene in B-cell lymphomas is the B-cell lymphoma 6 transcriptional repressor, and persistent expression results in malignant transformation of germinal center B cells[10]. BCL6-deficient animals lack GCs and are incapable of producing high affinity antibodies. Repression in macrophages would therefore be predicted to induce excessive cytokine release and gave an explanation for the inflammation observed. The mice also showed a severe inflammatory reaction marked by an eosinophilic infiltration of the heart, resulting in early death[11].

The BCL-6 gene is a 92- to 98-kD nuclear phosphoprotein which located on chromosome 3q27[12]. The BCL6 protein includes three conserved domains that are crucial to its functionality-the N-terminal BTB/POZ domain, a central RD2 region, a C-terminal zinc finger domain[2]. The protein’s N-terminal BTB-POZ and RD2 domains are in charge of attracting histone deacetylases and corepressor molecules, respectively[13]. Highly conserved[14] from Drosophila to Homo sapiens, the BTB domain is present in a broad and diverse family of proteins. BTB domains are the 28th most prevalent motif in the human proteome (29) and were discovered in 113 proteins. They are essential for development, homeostasis, and neoplasia[15]. The development and performance of follicular helper T cells and other helper T cell subsets were unaffected by the deletion of the BTB domain’s function[16]. Some viral and cellular proteins with the BTB domain are also found, albeit they lack any clear DNA binding motifs and their function may not be related to transcription[17].

Non-Hodgkin lymphomas (NHL) are a group of extremely diverse malignancies that primarily develop from germinal center (GC) B cells. Chromosome translocations and BCL-6 gene mutations are the most frequent genetic abnormalities that cause DLBCL[18]. The most frequent chromosomal aberration found in diffuse large B-cell lymphoma—a cancer marked by genetic heterogeneity and a wide variation in clinical outcome—is a BCL6 gene rearrangement[19]. Diffuse large B-cell lymphoma (DLBCL) is the most prevalent variety of lymphoma, which accounts for around one-third of all occurrences globally,[20][21][22][23] these are aggressive and it is a diverse group of illnesses with varying prognoses that are differently characterized by clinical aspects[20].About 10 to 20% of diffuse large B-cell lymphoma (DLBCL) individuals remained “unclassified” after gene-expression profiling differentiated the activated B-cell-like (ABC) and germinal-center B-cell-like (GCB) subtypes[24]. Patients typically appear with a quickly spreading tumor mass in one or more nodal or extranodal locations[22]. Large B-cell lymphomas, also known as “double hit” (DHL) or “triple hit” (THL) lymphomas when they have BCL2 and/or BCL6 rearrangements along with MYC, these types are biologically extra aggresive[25].

Chromosome translocations affecting the BCL6 locus at band 3q27 in tumours are the most prevalent and identifiable genetic aberration linked to DLBCL. These mutations prohibit BCL6 from connecting to its own promoter, which breaks down the protein’s negative autoregulatory circuit[26]. There was almost similar frequency of IgH and non-IgH partners involved in the translocated cases, and there was no relationship between the amounts of BCL6 mRNA and protein expression[27]. The mutation process is allowed with non-Ig promoters, despite the fact that the alterations are connected to transcription initiation. The genes for c-MYC, S14, or a-fetoprotein (AFP) did not show any notable alterations, however BCL-6 was shown to be substantially mutated in a significant number of memory B cells in healthy people[28].

In this study, we performed on the 20 ns simulation to compare the behavior of the wild type and three mutant proteins. By looking into the structural dynamics of the Q8C, R67C, and N84C mutations within the Bcl-6 protein, we are able to shed light on their possible contribution to the emergence of cancer. The simulation parameters and analyses offered here provide a foundation for further research into the molecular mechanisms behind the effects of these mutations on cancer growth. Our findings emphasizes the significance of taking into account the structural dynamics of certain mutations within the Bcl-6 protein in order to comprehend their functional effects on cancer. The mutant structures’ increased stability and altered flexibility point to possible ramifications for apoptotic signaling and cell survival. These discoveries shed important light on the molecular underpinnings of cancer growth and may aid in the development of tailored therapeutics to reestablish healthy apoptotic control. To confirm these findings and investigate the therapeutic potential of focusing on these particular mutations within this crucial protein family in cancer treatment, additional experimental research are required.

Calculations based on root-mean-square deviation (RMSD) were used to evaluate the overall structural stability and changes in comparison to the initial structure. The flexibility and dynamics of specific residues were also shown using root-mean-square fluctuation (RMSF) analysis, and the simulation’s compactness of the proteins was assessed using the radius of gyration (Rg). According to the RMSD analysis, our findings showed that the mutant structures had distinct conformational differences from the wild-type. The improved stability and compactness of the mutant structures indicated possible effects on protein function. The RMSF study brought attention to changes in a few residues’ flexibility, suggesting a potential impact on regional dynamics and functional areas. By interfering with apoptotic signaling pathways and encouraging cell survival, these dynamic changes may contribute to the onset and spread of cancer.

## 2. Methods and Materials

### 2.1. Structural Preparation

The primary sequence of the BCL6 BTB domain protein of the model organism Homo sapiens (Human) was retrived from the UniProtKB database in FASTA file format with the UniProtKB identifier P41182 consist of 123 amino acid long sequence. The physicochemical parameters of the protein BCL6 BTB domain were obtained using Expasy’s Prot param server. Additionally, we looked at the three BCL6 BTB domain mutant Q8C, R67C, N84C protein produced by the UCSF Chimera [29].

### 2.2. Molecular dynamics (MD) simulation

Molecular dynamics (MD) simulation has matured into a technique that can be effectively used to understand structure-to-function relationships in macromolecular systems [30]. It can predict the motion of each atom in a protein or other molecular system using a simplified model of the physics governing interatomic interactions. Molecular dynamics simulation can capture a variety of significant biomolecular phenomena, such as conformational changes, ligand association, protein folding, and precise atom positions at femtosecond temporal resolution [31]. Here, the wild type and three BCL6 BTB domain protein mutants were studied in molecular dynamics simulation to learn more about how they behave in trajectory analysis. The molecular dynamics simulation investigation was carried out according to the protocol outlined by Paul et al. [32].

A 20 ns (20000 ps) molecular dynamics simulation was performed to evaluate the stability and consistency of the predicted structure of the BCL6 BTB domain protein. This study was carried out using GROMACS 5.2 (2020.1) [33] within a Google Colab framework. The simulation was conducted to compare the trajectory data of the mutatant structures and to predict the similarities shared between the wild-type and mutatant structures trajectory analysis.

Within the Google Colab framework, we carried out the simulation by following a series of steps, where each step consists of a command, and where we merely executed the command. Before beginning the simulation, we first connected to the GPU and then installed GROMACS 2020.6 version for the purpose of preparing our MD system and carrying out our MD simulations, biopython for the purpose of managing PDB files, and py3Dmol for the purpose of depicting the protein structure. Then, we mounted our google drive to store the data in the drive. Following mounting, the present working directory was selected in the drive and then we imported the pdb files using biopython. To set up our MD simulation, we followed four basic steps named cleaning the atomic inputs, atoms parameterizing, protein solvation, and neutralization. In the cleaning step, we used ‘grep’ to remove the water molecules. In the parameterization step these proteins are parameterized using the AMBER99SB-ILDN force field [34], an improved version of Amber ff99SB force field which has limitations to the four residues (isoleucine, leucine, aspartate, and asparagine). AMBER99SB-ILDN force field is frequently used in MD simulations and its parameters accurately depict the dynamics and flexibility of folded proteins.

The AMBER99SB-ILDN force field[34], an upgraded version of the Amber ff99SB force field, is used to parameterize these proteins during the stage of parameterization. In MD simulations, the AMBER99SB-ILDN force field is commonly employed because the parameters of this force field accurately describe the dynamics and flexibility of folded proteins. Additionally, a universal equilibrium 3-point solvent model called the simple point charge (SPC) water model spc216 was used to solve the structures of the wild-type and three mutant proteins. The projected model was then enclosed in a 1.50 nm cube of simulation box and solvated with an SPC water model. The system was neutralized with 4Na+ ions while it was being solvated. Periodic boundary conditions were applied in all directions to eliminate edge effects.

The energy minimization of the system involved the use of the structures of the wild-type protein and three mutants. The steepest descent algorithm was employed, with a maximum step size of 50,000 and 1000 kJ mol-1 nm-1 tolerance. After system minimization, equilibration was performed for 100 ps under both isothermal-isobaric ensemble (NPT) and canonical ensemble (NVT). The Particle Mesh Ewald (PME) [35] summation was used for establishing long-range electrostatic interactions with a PME order. The Linear Constraint Solver (LINCS) algorithm was used to constrain the hydrogen-containing bonds [36]. Additionally, the SETTLE algorithm was employed to constrain the geometry of water molecules [37]. The Parrinello-Rahman method [38] controlled a constant amount of pressure at 1 atm (1.01325 bar), while the V-rescale weak coupling method controlled the temperature at 300K. The LINCS algorithm [36] and a 2fs (fs) integration stage were used for the 20 ns MD run output with no constraints. Additionally, the PME method [35] was used for the Coulombic and Lennard-Jones interactions [39] in this study. During the simulation we were interrupted several times then we used the appending code to run the simulation from where it is exactly interrupted. Finally, we concatenated all the xtc files to analyze the fundamental and essential dynamics.

### 2.3. Fundamental Molecular Dynamics Analysis

In order to assess internal fluctuation in distinct complexes, the root mean-square fluctuation (RMSF) of one chain was contrasted with that of the counter chain within the same complex[29]. It is possible to determine the motions of amino acids using the RMSF trajectory data from molecular dynamics simulation[32]. The protein’s compactness and overall structural changes during the simulation were assessed using the radius of gyration method. Calculating Rg is one of the most crucial measures that is usually used to estimate the structural activity of a macromolecule[40]. To analyze the fundamental dynamics, we used the google co-Lab protocol for four trajectory analysis. After successfully connecting to the GPU, we moved on to this part of installation of python, matplotlib, mdtraj, nglview, cython, python, pytraj, tsplot, and gnuplot-x11, and biotitle by making use of python installation packages. Following installation processes, we loaded the all xtc files and solv.ions files to evaluate the rmsd, rmsf and rg values for the wild type as well as three mutant proteins.

### 2.4. Essential Molecular Dynamics Analysis

The MDTraj Python package was also used to group the MD simulation trajectories of three BCL6 BTB domain protein mutations using RMSD distance matrices and hierarchical clustering approaches. A dendrogram with RMSD average linkage hierarchical clustering is produced by computing each pairwise RMSDs between conformations. Principal component analysis (PCA), is a frequently used technique for extracting functionally significant collective motions from a molecular dynamics (MD) trajectory. Using the previously mentioned techniques, we conducted an investigation of the essential dynamic [29][41][42]. PCA is the process of calculating the principal components and utilizing them to alter the data underlying structure, frequently using just the top few and ignoring the rest[42]. PCA is a technique for finding patterns in high-dimensional data and visually highlighting them to show their similarities and contrasts. Solvent accessibility (SASA), a crucial aspect of proteins for affecting their folding and stability, was taken into consideration to better understand three BCL6 BTB domain protein mutations correlated dynamics[43]. In our analytical section, SASA analysis identified the solvent-exposed region, which may reduce the protein’s solubility and affect protein-protein interactions[44][45]. To analyze the essential molecular dynamics, we also used the google colab protocol.

## 3. Results and Discussion

### 3.1 structural Preparation

We retrieved the whole structure of the wild-type BCL6 BTB Domain from the UniProtKB database. Using the Chimaera programme, three point mutation was inserted into the wild-type structure of the BCL6 BTB Domain by switching the residues Q8 to C8 (Glutamine to Cystein), R67 to C67 (Arginine to Cystein), and N84 to C84 (Asparagine to Cystein). The physiochemical parameters of these proteins were calculated using Expasy’s Prot param server where we used the amino acid sequence as input. The result (**Table.1)** shows that total number of atoms, molecular weight, theoretical PI, instability index and GRAVY changes followed by the mutations.

**Table-1:** Physiochemical characteristics of the 5N20 and mutants.

| Characters | Values |  |  |  |
| --- | --- | --- | --- | --- |
|  | WT | Q8C | R67C | N84C |
| **Number of amino acids** | 123 | 123 | 123 | 230 |
| **Total number of atoms** | 2008 | 2002 | 1996 | 2005 |
| **Molecular weight (Da)** | 14198.51 | 14173.52 | 14145.46 | 14187.54 |
| **Theoretical pI** | 8.02 | 7.94 | 7.01 | 7.94 |
| **Extinction coefficients (M<sup>-1</sup> cm<sup>-1</sup>) at 280 nm in water, assuming all pairs of Cys residues form cystines** | 4595 | 4595 | 4595 | 4595 |
| **Extinction coefficients (M<sup>-1</sup> cm<sup>-1</sup>) at 280 nm in water, assuming all Cys residues are reduced** | 4470 | 4470 | 4470 | 4470 |
| **Instability index (II)** | 54.29 | 55.38 | 50.64 | 50.72 |
| **Aliphatic index** | 105.37 | 105.37 | 105.37 | 105.37 |
| **Grand average of hydropathicity (GRAVY)** | 0.053 | 0.102 | 0.110 | 0.102 |

### 3.2 Fundamental Dynamics Analysis

The root mean square deviation (RMSD) is a widely used metric in analyzing molecular dynamics (MD) trajectories. Its primary purpose is to determine the equilibration period and assess the quality of biomolecular simulations. That is, the protein’s stability in relation to its conformation can be assessed by analyzing the deviations that occur during its simulation. A protein structure is considered to be more stable when the degree of deviation is smaller. Our study computed the RMSD value **(Fig. 1A)** for the C-alpha backbone over a 20-nanosecond simulation period to assess the stability of the wild-type, Q8C, R67C, and N84C systems.

**Figure-01:**
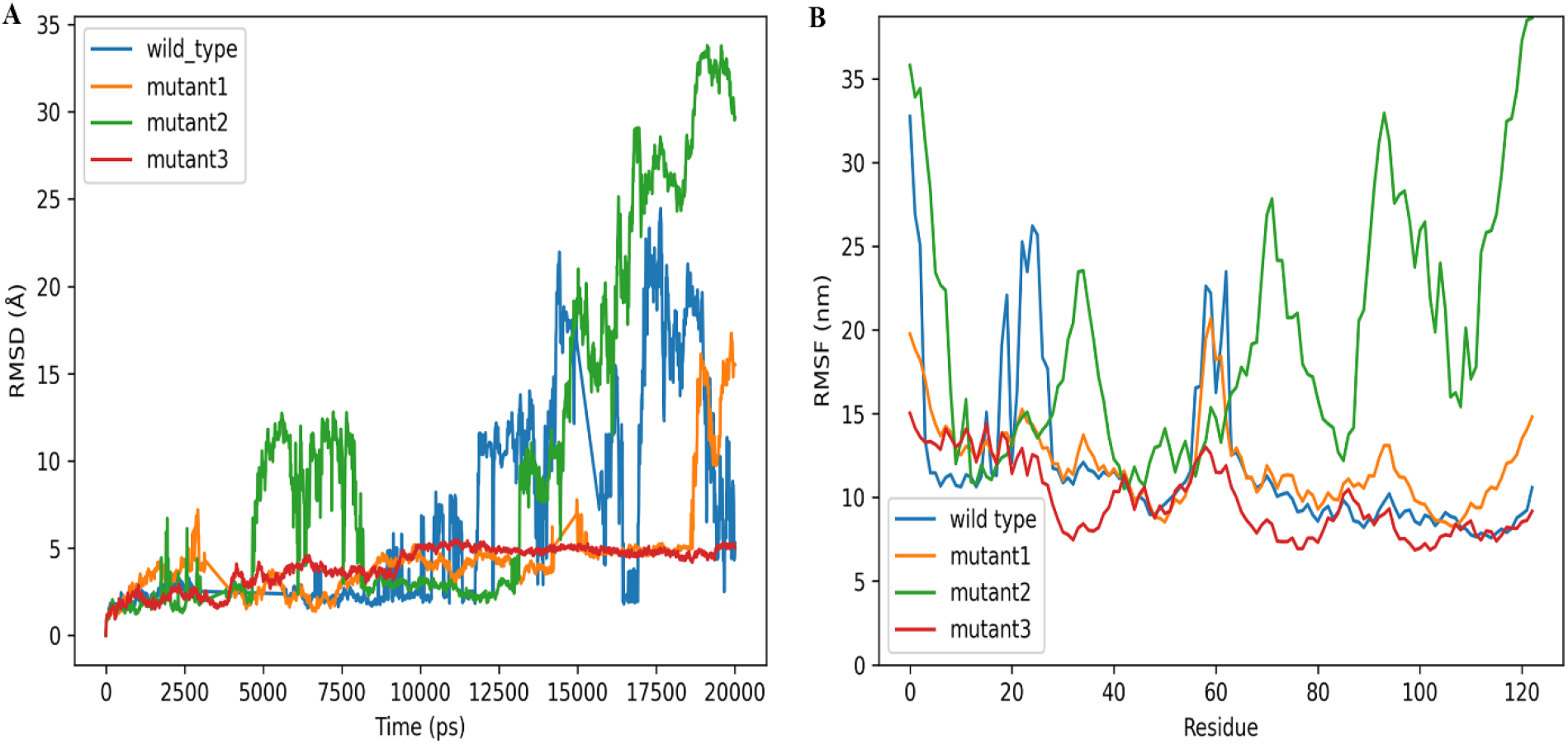
**(A)** Root mean square deviation (RMSD) plot of wildtype and mutant (Q8C, R67C and N84C) BCL 6 BTB domain proteins. Plot shows the compactness index for the WT, Q8C, R67C and N84C systems over 20 ns. **(B)**RMSF graphic illustrating the degree of structural flexibility and fluctuation in the wild-type (blue), mutant 1 (green), mutant 2 (orange), and mutant 3 (red) proteins for the BCL 6 BTB domain.

The RMSD simulations show that the wild-type system and the three mutations were in equilibrium up to 4800 ps. Then the fluctuation of mutation 2 was visible from 4800 ps to 8000 ps during the simulation. After that, the wild-type and all three mutants were compatible up to 13000 ps. A massive fluctuation occurs in mutant 2 after 13000 ps. Here, the range of RMSD values for mutant 2 was 0.2– 3.4 nm. The wild type also fluctuated from 9000 ps. It had a maximum fluctuation value of 2.5nm. In contrast, mutant 1 and mutant 3 had very low fluctuations. Therefore, mutant 2 showed higher RMSD variation values than wild-type, mutant 1, and mutant 3.

The RMSF is an averaged estimation of a mutant complex by which we forecast the displacement of a particular atom, or cluster of atoms, with regard to the wild-type structure. An RMSF plot **(Fig. 1B)** of alpha carbon atoms was used to evaluate the dynamic behavior of residues and protein flexibility. In comparison to the WT, Mutant-1 (Q8C), and Mutant-3 (N84C) systems, RMSF analysis clearly demonstrated more flexibility for the Mutant-2 (R67C) mutant system, which has three extremely flexible areas (residues 32–40, 72–84, and 89–110 where the deviation range were 16-24 nm,17-28nm and 12-33nm). Wild type structure is less compact than Mutant-1 (Q8C), and Mutant-3 (N84C) and WT has two flexible areas (residues 17-20 and 20-25 where the deviation range were 15-23 nm and 16-26nm). We can see from the RMSF value that the Mutant-1 (Q8C), and Mutant-3 (N84C) structure is more compact and has fewer deviations in its trajectory, which represents its stability.

The radius of gyration (RG) analysis indicates the proteins compactness. A protein has a tendency not to fold readily if it is extremely compact. When it is less stable that time the more it fluctuates. A highly unstable system is indicated by a larger Rg value, whereas a stable and equilibrium system is implied by a lower Rg numbe. In the Rg trajectory **(Fig. 2)** there is sudden change in the radius of the complex which show how the compounds interference with one another structure. At first R67C is fluctuate 5000 to 8000 ps where the Rg value is 18nm. The highest Rg value for the R67C, which is started in 13500ps and its rg value 38nm. In this figure we show that N84C is more stable than others and R67C is less stable as it is more fluctuates.

**Figure-2:**
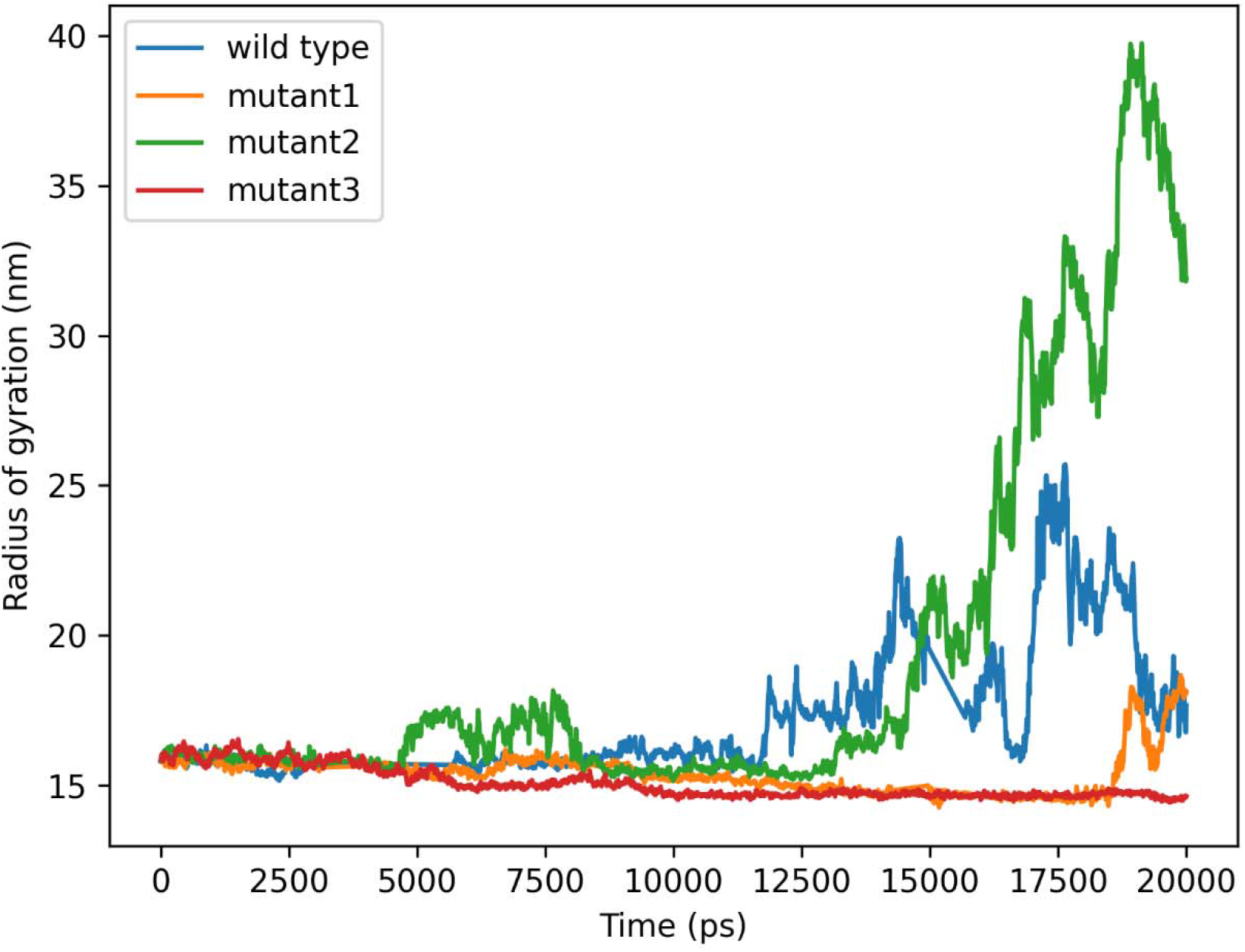
Analysis of the radius of gyration (Rg) for wild-type and mutant (Q8C, R67C, and N84C) BCL 6 BTB domain proteins during 20 ns MD (molecular dynamic) simulations period. For the BCL 6 BTB domain MD, molecular dynamics simulation, the wild-type protein is shown (blue), mutant 1 (green), mutant 2 (orange), and mutant 3 (red) are represented in the proteins. The radius of gyration is a monitor used in MD simulation to track the generation of new structures. According to Figure 2, the gyration radius profile of the wild-type protein dramatically dropped, whereas the mutant-2 protein structure showed the largest peaks.

### 3.3 Essential Analysis

In this study, we executed the RMSD average linkage hierarchical clustering, Cartesian coordinate principal component analysis, pairwise distance principal component analysis, Solvent Accessible Surface Area (SASA), and SASA autocorrelation of 20 ns simulation trajectory in order to gain an understanding of the conformational behavior and to evaluate the wild-type protein in comparison to three distinct mutations.

After running MD simulations of proteins with the wild-type and three different mutations, we used a hierarchical clustering technique based on root-mean-squared deviation (RMSD) to create a trajectory dendrogram **(Fig.3)** for 20 ns simulation. The hierarchical structure and degree of similarity between the conformations are displayed by the dendrogram. The four different conformations of the Wild-type protein are shown in orange, green, red, and violet in **Fig. 3A**. Mutant 1 trajectory (**Fig. 3B)** demonstrates that it has two distinct conformations, colored orange and green, respectively. For mutant 2(**Fig. 3C)**, we see only one conformation. Moreover, three different conformations have been demonstrated (**Fig. 3D)** for mutant 3. These findings point to structural modifications as a result of mutation.

**Figure-3:**
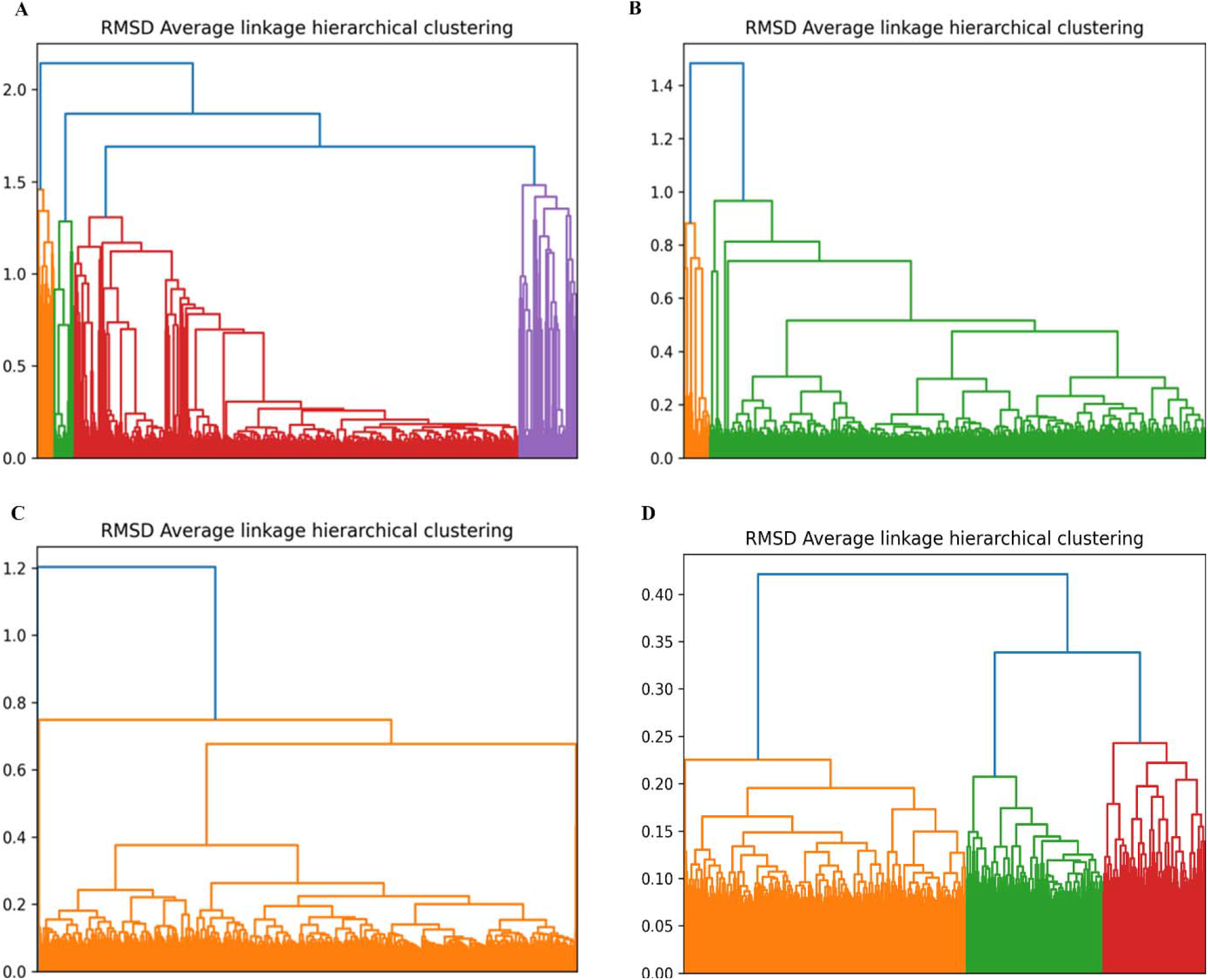
Crucial dynamics analysis of the structures of (A) wild-type, (B) mutant-1 (Q8C), (C) mutant-2(R67C), and (D) mutant-3 (N84C). It depicts both structures’ clustering dendrograms over the course of the whole 20 ns simulation period. By using RMSD hierarchical clustering, the hierarchical distribution of each cluster is displayed.

Principal component analysis (PCA) has become one of the most popular methods for analyzing the movement of proteins within this framework. Here, PCA analysis of the collective motions and conformational changes in the BCL-6 BTB domain protein wild-type, mutant-1 (Q8C), mutant-2 (R67C), and mutant-3 (N84C) structures. The pairwise distance PCA is computed by analyzing the precise location of the atoms over time. In contrast, the Cartesian coordinate PCA analysis captures a portion of the dominant overall motion. RMSD hierarchical clustering was represented by color-coded clusters against a fixed time frame and color distribution. Both Cartesian coordinate PCA and pairwise distance PCA simulations spanned 20ns from the initial to the final stage.

The Cartesian coordinate PCA diagrams in figures 1A,1B,1C, and 1D depict the entire 20ns of the entire simulation period for the wild type (A) and all three mutated structures (B, C, D) on a 2D graph. **In Figure 4A**, we observed that initial clusters resided between the first and fourth coordinates, whereas in the final stage, they were located in the second and third quadrants. **In Figure 4B**, the primary clusters were located in the fourth quadrant, and their final appearance was between the second and third quadrants. The initial clusters were observed to be located within the range of the first and fourth quadrant in **Figure 4C**. During the final phase, the subjects were positioned in the second and third quadrant. In **Figure 4D**, We also visualized initial clusters located between 1st and 4th quadrant. Their final appearance was between 2nd and 3rd quadrant as well.

**Figure-4:**
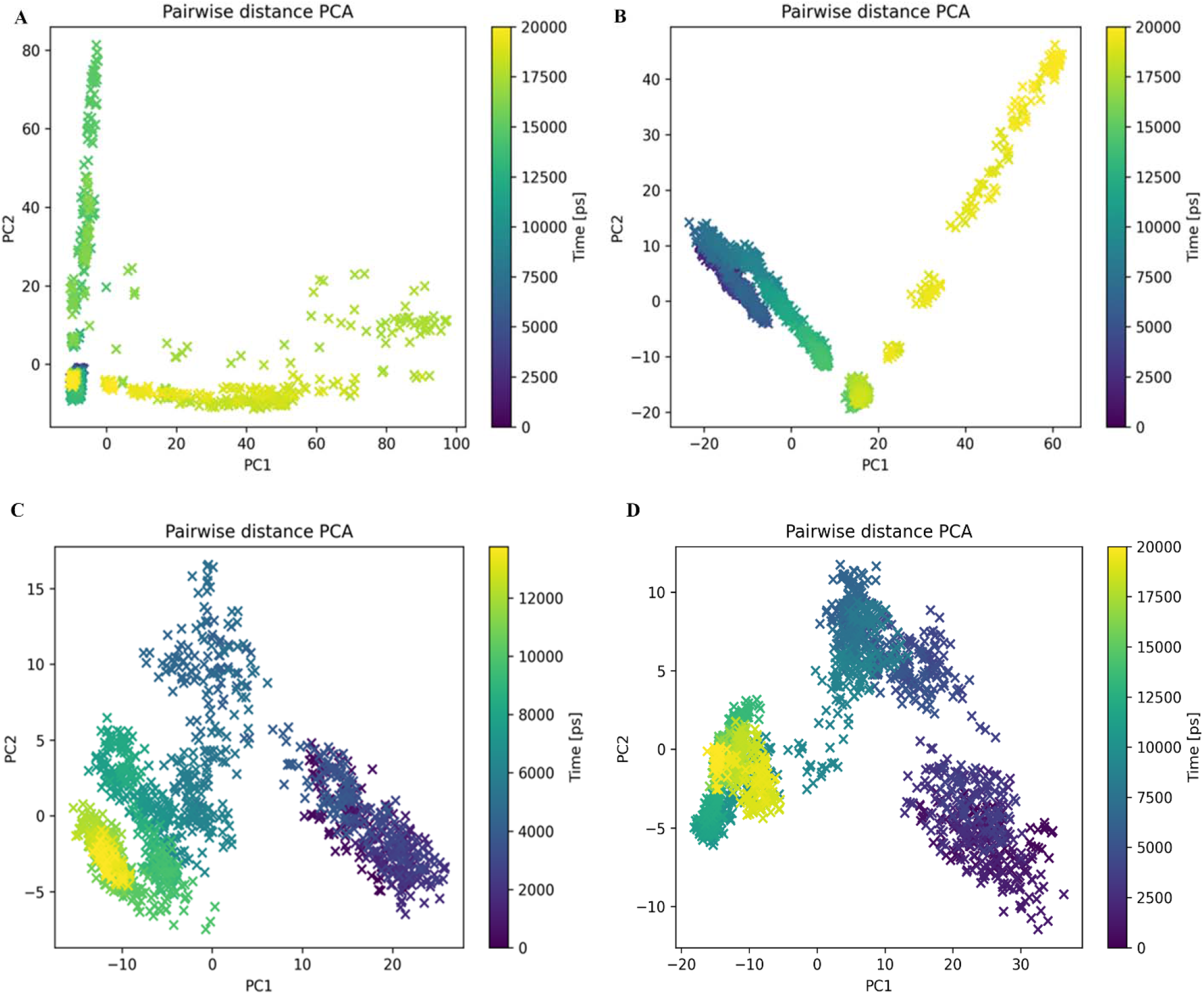
Visualizing pairwise distance (PCA) principal component analysis of the wild-type and mutant structure of entire 20ns of the simulation period. (A) Visualization of pairwise distance PCA of the wild type. (B) Visualization of pairwise distance PCA of mutant-1 (Q8C). (C) Visualization of pairwise distance PCA of mutant-2(R67C). (D) Visualization of pairwise distance PCA of mutant-3 (N84C). Different conformation states are depicted by colors ranging from dark blue to pale yellow, while transition stages are symbolized by the color purple.

In the pairwise distance PCA, the initial cluster in the wild type **(Fig. 5A)** was in the third quadrant and the final cluster was in the third and fourth quadrants. In mutant 1**(Fig. 5B)**, we observed the initial cluster between the second and third quadrants and the final appearance was in the first and fourth quadrants. The initial cluster of mutant 2 was located in the first and fourth quadrants **(Fig. 5C)**, while the final cluster was found in the third quadrant. Finally, in the third mutant **(Fig. 5D)**, the initial cluster was located in the first and fourth quadrants, while the final cluster was observed in the second and third quadrants.

**Figure-5:**
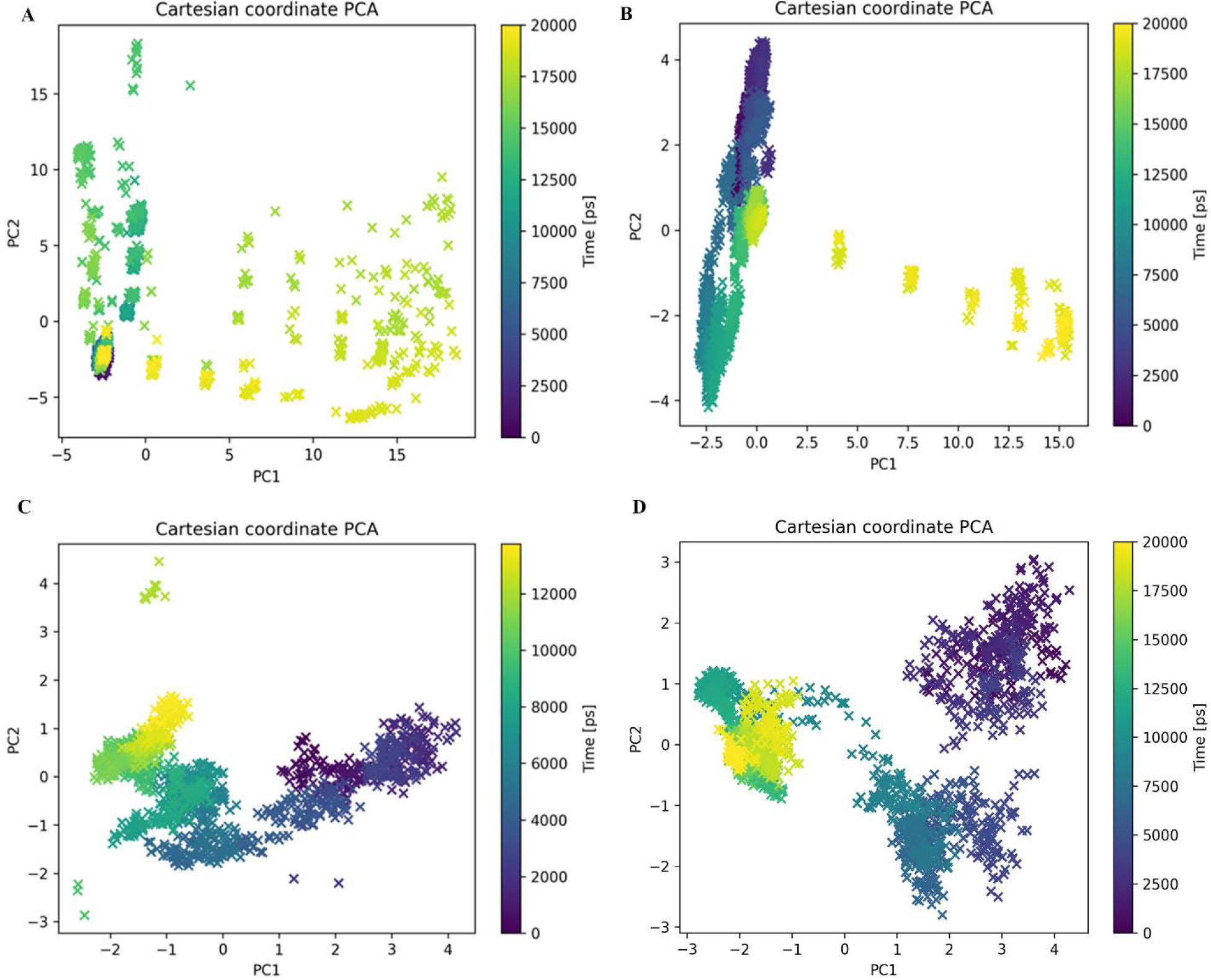
Visualizing Cartesian coordinate (PCA) principal component analysis of the wild-type and mutant structure of entire 20ns of the simulation period. (A) Visualization of Cartesian coordinate PCA of the wild type. (B) Visualization of Cartesian coordinate PCA of mutant-1 (Q8C). (C) Visualization of Cartesian coordinate PCA of mutant-2(R67C). (D) Visualization of Cartesian coordinate PCA of mutant-3 (N84C). Different conformation states are depicted by colors ranging from dark blue to pale yellow, while transition stages are symbolized by the color purple.

A biomolecule surface area that is accessible to a solvent is known as the solvent accessible surface area (SASA). The increase in SASA value indicates relative expansion. We calculated SASA for both the wild type and the mutant proteins, which is essential for hydrophobic core region analysis to accurately comprehend the protein’s stability, binding interaction, and folding pattern. The wild type plot **(Fig. 6A)** showed value of 73-80 nm^2^, where the highest peak value of about 81 nm^2^ at approximately 18 ns. The mutant 1(Q8C) trajectory **(Fig. 6B)** showed that the average SASA value is 72-77 nm^2^, containing the highest value is 77 nm^2^. The SASA value for the mutant 2 (R67C) and the mutant 3 (N84C) is 72-76nm^2^ and 73-77 nm^2^ respectively**(Fig. 6C-6D)**.

**Figure-6:**
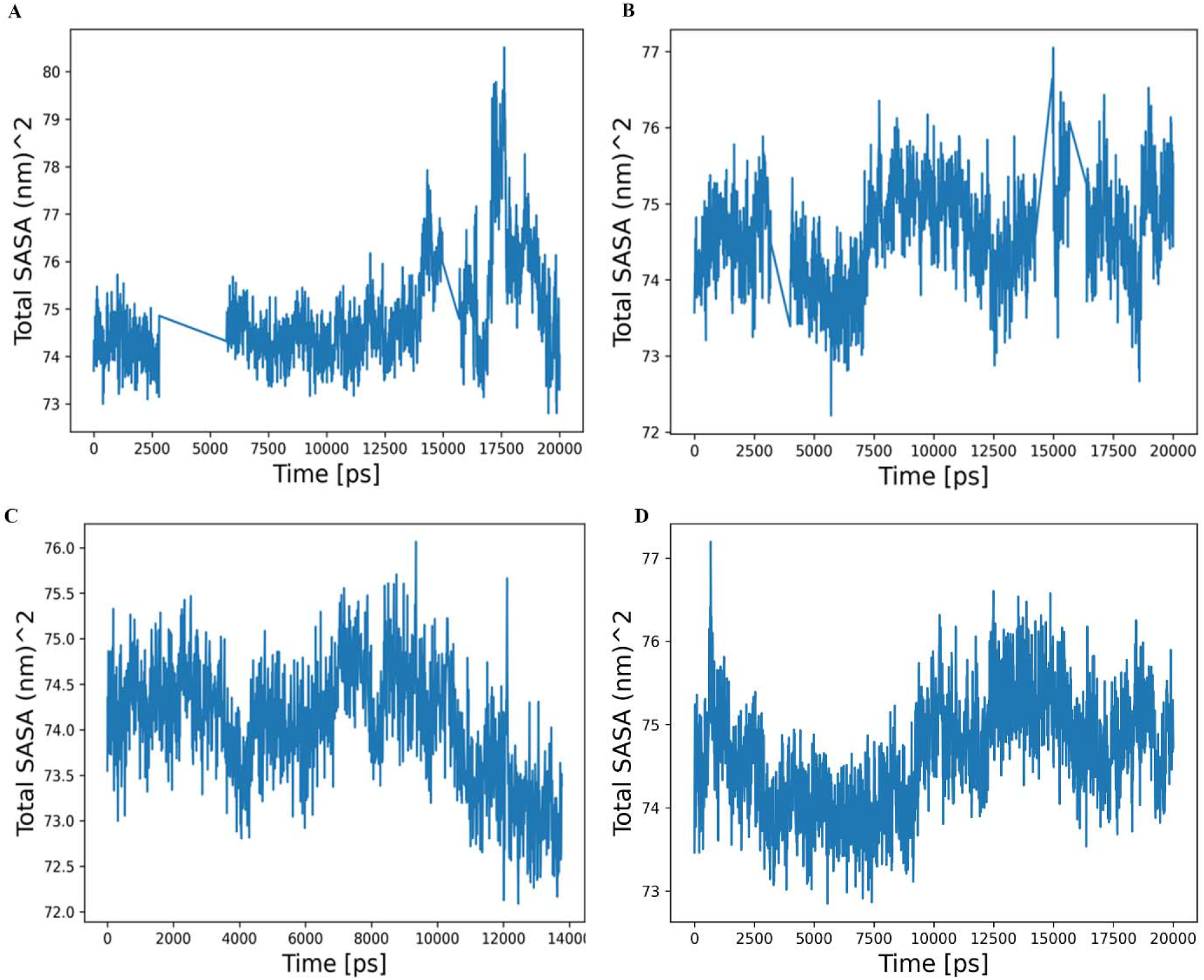
Solvent accessible surface area (SASA) analysis of wild and mutant structure of BCL 6 BTB domain proteins during 20 ns MD (molecular dynamic) simulations period. Blue, green, orange and red lines indicate wild type, Q8C, R67C and N84C proteins respectively.

SASA autocorrelation is a term used in the science of molecular dynamics to describe the analysis of correlation in the time-dependent fluctuations of a molecule’s solvent accessible surface area during a simulation. It reveals light on the chemical process and spatial behavior of the molecule’s surface exposure.

In figure 7, SASA autocorrelation was displayed. The SASA autocorrelation of the **(Fig. 7A)** wild-type protein (A) showed a steady value of around 0.8 nm^2^ for the first 1-10 ns, followed by a consistent decrease until two peaks were observed at 0.3 nm^2^ (1.1-1.3 ns) and 0.2 nm^2^ (1.4-1.5 ns). The mutant protein **(Fig. 7B)** Q8C showed a similar trend, with a steady value followed by a consistent decrease until two peaks were observed at 0.1 nm^2^ (8-10 ns) and 0 nm^2^ (11-12 ns). The mutant proteins **(Fig. 7C)** R67C and **(Fig. 7D)** N84C showed a steady value of around 0.6 nm^2^ for the first 1-10 ns, followed by a consistent decrease until a peak was observed at 0.1 nm^2^ (8-10 ns) for R67C and -0.1 nm^2^ (8-10 ns) for N84C.

**Figure-7:**
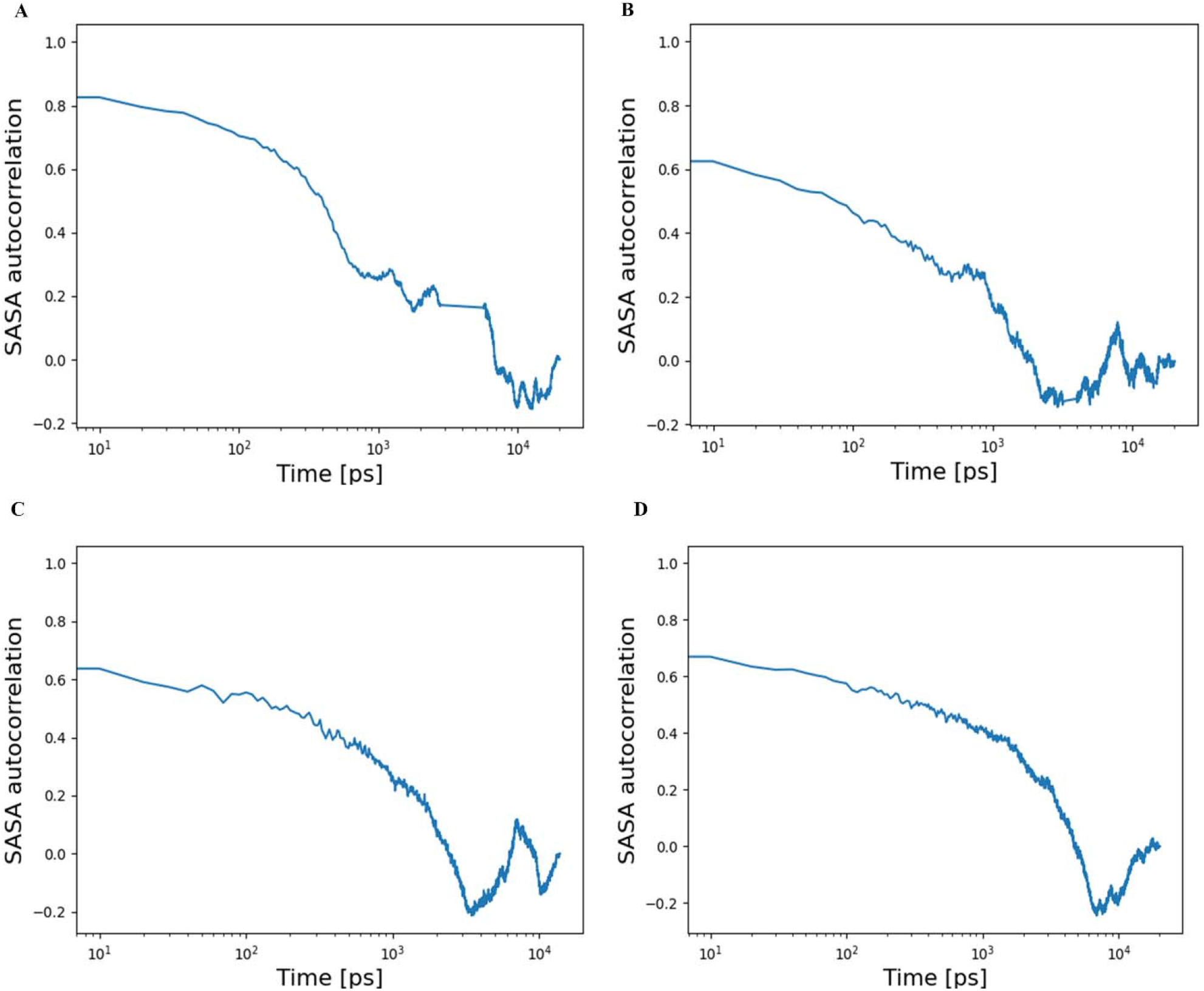
SASA autocorrelation is demonstrated. The wild type (7A) variant maintained a constant value of around 0.8 nm2 between 0.10-1 ns, a continuous decline until two peaks of approximately 0.3 nm2 (at 1.1 and 1.3 ns) and 0.2 nm2 (at 1.4 and 1.5 ns) were noticed between 1-10 ns. In (7B), a steady decline was noticed until two maxima, averaging around 0.2 nm2 each (at 8–10 ns and 11–12 ns), were noticed between 1–12 ns. Similar trend was seen in mutant structure (7C and 7D), where a steady decline was noticed until one peak, measuring around 0.2 nm2 (at 8–10 ns), was detected.

## 4. Conclusion

Computational analysis has proven to be an invaluable tool for studying the structure of proteins, their conformational properties, dynamics and surface properties. It plays a significant role in the characterization of protein mutations at the atomic level. The molecular dynamics (MD) simulation is a very well-established computational analysis used to explore the relationship between protein structure and function. The main purpose of this study was to apply computational methods to evaluate the effects of structural changes on BCL6 BTB domain protein mutations. So, we conducted an analysis of 20 nanoseconds (20,000 picoseconds) molecular dynamics simulation of the BCL6 BTB domain protein and its three mutant Q8C, R67C and N84C. The conformational changes of all protein structures are monitored using trajectory analysis techniques such as RMSD, RMSF, Rg, PCA, clustering analysis, SASA, and SASA autocorrelation.

According to the results we obtained, mutant 2 (R67C) caused a conformational change with a high RMSD value and exhibited a high degree of flexibility according to the RMSF value. This finding suggests that mutant 2 (R67C) exhibits greater protein instability than the wild-type BCL-6 BTB domain protein and the other two mutants. Moreover, the analysis of the Radius of gyration (Rg) suggests that mutant 3 (N84C) exhibits a higher compactness level than the remaining proteins. The principal component analysis (PCA) results also provide insights into the overall kinetic and conformational changes exhibited by the BCL6 BTB domain protein and its three mutant variants. Additionally, SASA analysis indicated that the BCL-6 BTB domain wild-type protein exhibits higher solvent accessibility. On the other hand, SASA autocorrelation calculates the temporal correlation of the solvent accessible surface area of proteins during a molecular dynamics (MD) simulation. However, the analysis of the structural dynamics of the Q8C, R67C, and N84C mutations reveals their impact on the growth of various immune system cells, such as germinal center B cells and CD4+ follicular helper T cells. The altered stability and flexibility of the mutant structures indicate the possibility of apoptotic signaling and cell survival. Furthermore, investigation of the structural dynamics of these mutants can provide insight into their potential role in cancer development. Additional experimental studies are needed to explore the therapeutic potential of targeting these specific mutations in cancer treatment.

## Acknowledgements

We would like to thank Shamrat Kumar Paul for giving us with simulation and analysis source code.

